# Integrating stereotypes and factual evidence in interpersonal communication

**DOI:** 10.1101/2023.05.23.540979

**Authors:** Saskia B. J. Koch, Anna Tyborowska, Hannah C. M. Niermann, Antonius H. N. Cillessen, Karin Roelofs, Jana Bašnáková, Ivan Toni, Arjen Stolk

## Abstract

Stereotypes can exert a powerful influence on our interactions with others, potentially leading to prejudice when factual evidence is ignored. Here, we identify neuroanatomical and developmental factors that influence the real-time integration of stereotypes and factual evidence during live social interactions. The study uses precisely quantified communicative exchanges in a longitudinal cohort of seventeen-year-olds followed since infancy, testing their ability to moderate stereotype tendencies toward children as contrary evidence accumulates. Our results indicate that the impact of stereotypes on communicative behavior is linked to individual variations in gray matter density and cortical thickness in the right anterior cingulate gyrus. In contrast, the ability to moderate stereotype tendencies is influenced by developmental exposure to social interactions during the initial years of life, beyond the effects of familial environment and later experiences. These findings pinpoint a key brain structure underlying stereotype tendencies and suggest that early-life social experiences have lasting consequences on how individuals integrate factual evidence in interpersonal communication.

## Introduction

Over the past century, social psychology has extensively studied the pervasive nature of stereotypes in human experience and society, offering compelling evidence of their impact on our initial impressions of others^1–3^. Stereotypes are generalized beliefs about groups organized along dimensions like warmth and competence, offering a powerful framework for making predictions about individuals^4–7^. Indeed, even the mere belief of interacting with a younger, less competent partner can lead to more emphatic communication with that individual^8–11^. However, overreliance on stereotypes can prove problematic if their influence is not adjusted to contrary evidence arising from social interactions. For example, an academic’s didactic tone may come across as patronizing when the young individual being spoken to turns out to be a peer rather than a student. Ignoring factual evidence in favor of these tendencies not only fuels prejudice but may also perpetuate systemic bias in society linked to stereotypes^12,13^.

The ability to deal with interaction-based evidence has been the main focus of constructivist approaches found in developmental psychology^14–17^. Throughout life, humans engage in joint activities with only partial knowledge, yet they consistently bridge interpersonal disparities through collaborative efforts to achieve mutual understanding^18–20^. This interactional capacity is thought to develop ontogenetically through social experiences necessitating the coordination of perspectives^21,22^. Empirical evidence corroborates this notion, indicating that children’s emerging appreciation of differing perspectives, as assessed through false-belief understanding, is influenced by their access to social interactions^23,24^. However, it remains unclear how stereotypes and interaction-based evidence combine to support interpersonal communication, as these phenomena are generally studied separately in their respective fields of psychology, often in isolation from interactive social settings^25^.

Here we sought to identify neuroanatomical and developmental factors that influence the real-time integration of stereotypes and factual evidence during live social interactions. Rather than relying on societal stereotypes associated with attributes like race and gender, we centered on the general expectation that younger individuals might need more explanatory guidance in communicative settings^26^. We employed a nonverbal communication game designed to differentiate between behavioral adjustments driven by participants’ stereotype beliefs about an interaction partner and those influenced by the dynamics of the interaction itself. Participants engaged in dyadic interactions on a digital game board, where they were challenged to generate novel nonverbal behaviors that could be mutually understood within the ongoing interaction^27,28^. This game design allowed precise measurement of communicative behavior and its dynamics across interactions while minimizing the influence of linguistic conventions. Additionally, participants were physically distanced from their interaction partners, allowing the substitution of expected partners (a child and an adult) with a single role-blind confederate. The presumed partners differed solely in terms of participants’ stereotype beliefs regarding their cognitive abilities, creating a conflict as participants balanced accumulating evidence from interactions (where both partners demonstrated equal competence) against pre-existing stereotype-based beliefs (where one partner was perceived as a less competent child). Adjustments driven by stereotype beliefs were quantified through participants’ spontaneous tendency to place greater emphasis on communicatively relevant aspects of behaviors directed at the presumed child partner. Adjustments influenced by the interaction dynamics were measured through participants’ convergence in communication toward the matched behaviors and understanding of the two presumed partners.

Our study builds upon two sets of observations regarding the neuroanatomical and developmental bases of stereotype beliefs and human interactional abilities, each of which provided specific predictions to disentangle their contributions in the current sample. First, given previous observations in neuropsychological patients^11,29–31^, we hypothesized that medial prefrontal brain structures, particularly in the right hemisphere^32,33^, play a role in leveraging stereotype beliefs when tailoring communication to an individual partner. Damage to this cortical region can distort judgments influenced by gender stereotypes and other social connotations^29–31^; notably, during performance of the same task, patients with medial prefrontal lesions fail to differentiate their communication between the presumed child and adult partner^11^. Second, prior developmental work has found that the amount of time children spend in daycare during their early years predicts how flexibly they adjust their communication to a partner at age 5, even after considering the influence of familial environment^10^. Humans are exceptional among existing hominids for experiencing early developmental exposure to unrelated conspecifics^34^, and it has been suggested that the initial years of life may represent a sensitive developmental period for social interaction^35–40^. Building on these considerations, we conducted a pre-registered analysis of a cohort of seventeen-year-olds who have been followed since infancy, in order to evaluate whether the consequences of social experiences acquired in a daycare environment persist through development to influence stereotype- or interaction-based communicative adjustments later in life^41^.

## Results

### Generation of Mutually Understood Communicative Behaviors

Ninety-five participants successfully completed the communication game after their seventeenth birthday (47 females; 17.2 ± 0.2 y of age). On each of forty consecutive trials, successful communication required participants to inform their interaction partner, a confederate, about the location of an object hidden in one of thirteen possible locations on a digital game board with a 3×3 grid layout (Fig. 1a). Only participants knew the object location, and only the partner could collect the object, leading participants to generate communicative behaviors of an avatar on the digital board that their partner could interpret to understand where the object was located. These circumstances drive dyads to achieve mutual understanding by aligning on a limited but idiosyncratic number of strategies from an open-ended set of possibilities^8^. Several features of the game challenged dyads toward achieving this understanding. First, the avatar could only be moved to the center of each of the nine grid squares, such that it could not be overlaid on the precise location of the object when a square contained multiple candidate locations. Importantly, even in trials where the object was located in a square with one possible location, participants needed to find a way to disambiguate the square containing the object from other squares for their partner, particularly squares that were visited while moving across the game board. Second, trials where the object was in a square with one, two, or three possible locations were pseudo-randomly intermixed such that there was an overall increase in difficulty of the game over the course of the experiment. Overall, dyads solved the trials well above chance (89.8 ± 11.7%, estimate of chance level: 1/13th, i.e. 7.7%), indicating that participants generated communicative behaviors that, despite their novelty, could be mutually understood.

**Figure 1.**
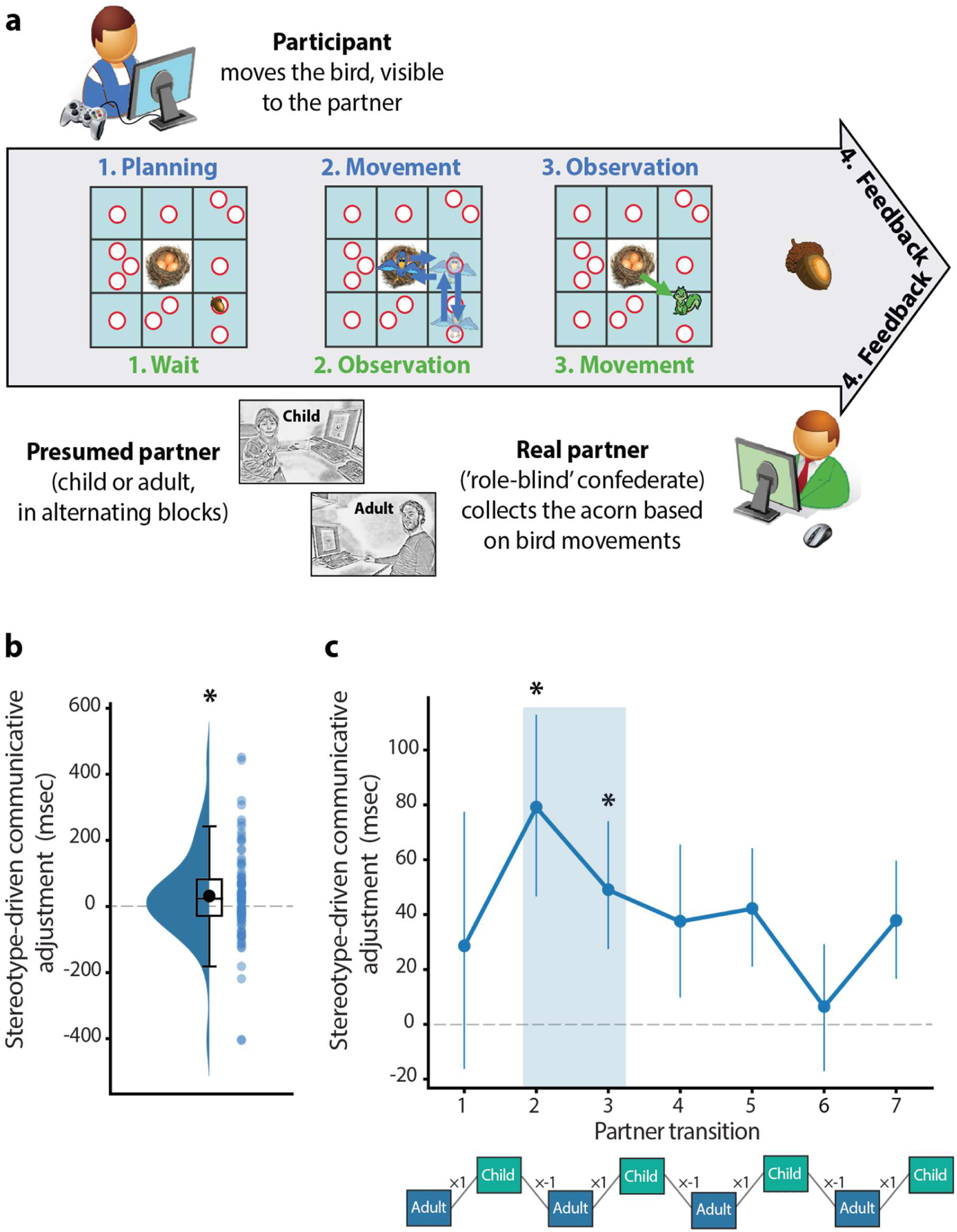
Nonverbal communication game and stereotype-driven adjustments. (**a**) In the game, participants had to inform their interaction partner about the location of an acorn on a digital game board. The acorn could be located in any of the thirteen white circles and its location was only known to the participant (phase 1). The participant could convey the acorn’s location to the partner by moving a bird avatar across the game board, visible to both players (phase 2). Starting at the nest (center square), the bird could only be moved to the center of each of the nine squares, which made it difficult for the avatar to be overlaid on the precise location of the acorn when a square contained multiple candidate locations. After the bird was returned to the nest, a confederate partner could move a squirrel avatar to the circle deemed to contain the acorn given the bird’s movements (phase 3). The squirrel movements were also visible to both players, allowing participants to interpret how their movements of the bird were understood by their partner, in addition to explicit feedback on the successful retrieval of the acorn at the end of the trial (phase 4). Feedback consisted of an acorn for successful trials, and an acorn with a superimposed red cross for unsuccessful trials. Importantly, participants were told they would be alternately interacting with an adult and a child partner. In reality, the confederate performed both roles, such that the two presumed partners differed only in terms of participants’ stereotype beliefs regarding their cognitive abilities. (**b**) Participants spent longer on communicatively relevant portions of the game board, i.e. squares containing the acorn, when producing behaviors directed at the presumed child partner. (**c**) A time-resolved analysis focused on partner transitions revealed that these stereotype-driven communicative adjustments were transiently prominent at the onset of interactions, including partner transitions 2 and 3, and gradually declined as interactions unfolded. Error bars indicate M ± SEM. Black asterisks denote statistically significant adjustments, *p* < 0.05, corrected for multiple comparisons. The experimental structure shown below the graph is one of two counterbalanced variants, starting with either the adult or child partner.

### Spontaneous Communicative Adjustment to Stereotype Beliefs

Previous research using the same task has shown that participants tend to generate subtle adjustments when believing to be communicating with a younger partner, resulting in longer time spent in squares containing the hidden object when moving across the game board^8–11^. This study exploits these communicative adjustments as a real-time quantitative marker of stereotype beliefs about an interaction partner, considering both the presence and the dynamics of stereotype-driven communicative adjustments. First, we replicated the presence of stereotype-driven adjustments in the current sample. These adjustments were driven exclusively by participants’ belief of interacting with a younger, less competent partner, as the behavior and understanding of the child and adult partner were matched by virtue of being produced by the same role-blind confederate. Across all trials, participants spent more time in the square containing the object when they believed they were communicating with the child partner (Fig. 1b, *M difference* = 32 ms, *t*_(94)_ = 2.31, *p* = 0.023, Cohen’s *d* = 0.24). These adjustments were specific to communicatively relevant portions of the game board, as other aspects of participants’ behavior were not affected by stereotype beliefs, including planning time and time spent in other locations of the game board (Fig. S1 and SI results). Moreover, the adjustments were not a consequence of differences in the confederate’s behavior or understanding, as the confederate showed similar task performance when performing the roles of the child and adult partner (Fig. S2 and SI results). Second, we performed a time-resolved analysis of stereotype-driven adjustments focused on partner transitions. This was achieved by computing the difference in average time spent in the object location between the eight consecutive experimental blocks of five trials, producing seven partner transitions (see experimental structure shown at the bottom of Fig. 1c). The sign of these partner transitions considered the expected direction of the adjustment, i.e. communicative adjustments were multiplied by +1 and -1 for adult-child and child-adult transitions, respectively. The resulting adjustment estimates were entered into a cluster-based statistical analysis correcting for multiple comparisons across transitions^42^. This analysis identified a cluster of positive adjustments spanning the second and third transition (Fig. 1C), suggesting that adjustments to stereotype beliefs were transiently prominent at the onset of interactions and gradually declined as interactions unfolded.

### Medial Prefrontal Brain Structures Leverage Stereotypes

Building on previous observations in prefrontal lesion patients^11,30^, we tested the hypothesis that right-hemispheric medial prefrontal brain structures play a role in leveraging stereotype beliefs when tailoring communication to an individual partner. We analyzed two computationally-independent in-vivo measures of neuroanatomical structure, based on voxel- and surface-based morphometry of T1-weighted MRI data^43,44^. These measures were used as predictors of stereotype-driven communicative adjustment in separate multiple linear regression analyses controlling for factors such as gender, scanner type, and salivary testosterone levels (a proxy of pubertal developmental status^45^). To accommodate variations in brain hemispheric volume among participants and improve sensitivity in detecting lateralized effects, we computed a voxelwise asymmetry index for each individual^43^. This index quantifies gray matter density in the right hemisphere relative to homotopic regions in the opposite hemisphere. A whole-brain voxel-based analysis revealed a correlation between communicative adjustment to stereotype beliefs and gray matter asymmetry within the anterior cingulate gyrus (ACCg; Fig. 2a, MNI_x,y,z_ = [3, 33, 24], *p*_FWE_ < 0.014, whole-brain corrected). Subsequent analyses focusing on hemispheric gray matter content extracted from this ACCg cluster revealed a positive correlation between stereotype-driven adjustment and ACCg volume in the right hemisphere (*ρ*_(65)_ = 0.516, *p* < 0.001), with no observed association in the left hemisphere (*ρ*_(65)_ = -0.130, *p* = 0.29). As seen in Fig. S3, this cortical region showed considerable neuroanatomical overlap with medial prefrontal brain tissue previously identified as crucial for tailoring interactions to stereotype beliefs about a partner^11^. Additionally, the impact of right ACCg on stereotype-driven adjustment proved statistically robust to replication in two additional datasets, including the same sample at age 14 and an independent sample performing the same task (Fig. S4). Notably, even after adjusting for right ACCg volume at age 14, the association between stereotype-driven adjustment at age 17 and right ACCg volume remained significant, underscoring the timeliness of this structural brain-behavior relationship (SI results). A surface-based analysis of cortical thickness corroborated these findings, revealing a positive association between stereotype-driven adjustment and cortical thickness in the right ACCg (*ρ*_(65)_ = 0.286, *p* = 0.019), based on the caudal anterior cingulate parcel from the Desikan-Killiany anatomical atlas^46^. Furthermore, adjustment to stereotype beliefs covaried with cortical thickness in a distributed right-hemispheric cortical network showing a degree of overlap with the theory-of-mind network^47–49^, including the superior frontal gyrus, superior temporal sulcus, temporoparietal junction, and precuneus (Fig. 2b, *p*_FWE_ < 0.05, whole-brain corrected). As discussed further below, these voxel- and surface-based neuroanatomical relationships remained statistically significant after controlling for variance accounted for by early social experiences (ACCg volume: *ρ*_(64)_ = 0.484, *p* < 0.001; and thickness: *ρ*_(64)_ = 0.362, *p* = 0.003).

**Figure 2.**
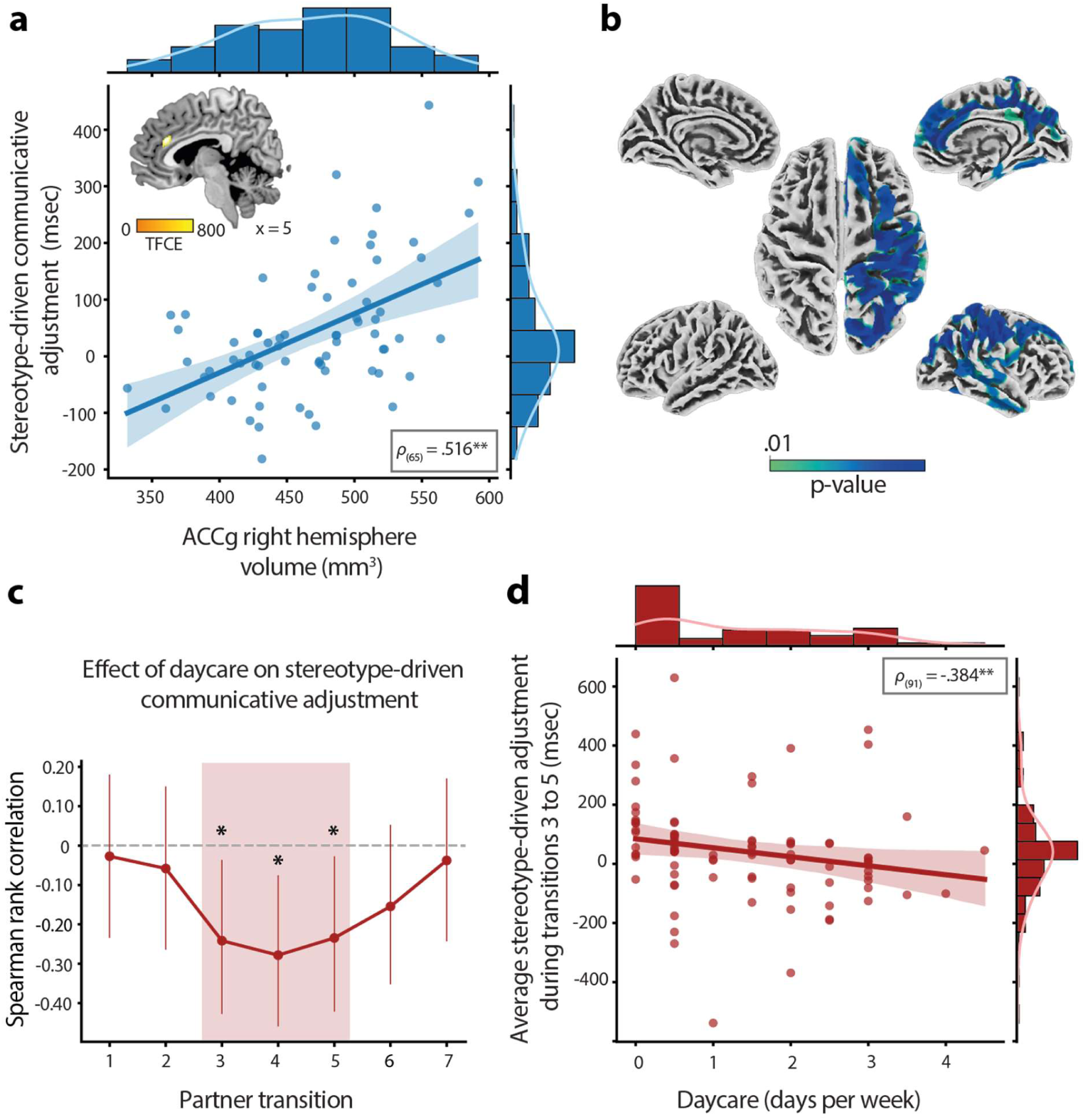
Neuroanatomical and developmental factors influencing stereotype- and interaction-driven adjustments. (**a**) Contributions from stereotypes correlated with gray matter density in the right anterior cingulate gyrus (ACCg). Statistical effects thresholded at *p*_FWE_ < 0.05 whole-brain corrected. Black asterisks denote *p* < 0.001. (**b**) Contributions from stereotypes also covaried with cortical thickness in ACCg as well as in a distributed right-hemispheric cortical network showing a degree of overlap with the theory-of-mind network. Statistical effects thresholded at *p*_FWE_ < 0.01. (**c**) Diminishing contributions from stereotypes during partner transitions 3 through 5 correlated with time spent in daycare during the first few years of life. As seen in Fig. 1C, this interaction-driven effect coincides with a reduction in stereotype-driven adjustments across the group, with individuals who spent more time in daycare showing a faster rate of communicative convergence toward the matched understanding of both partners. Error bars indicate 95% confidence intervals. (**d**) Scatterplot of the inverse relationship between daycare attendance and stereotype-driven communicative adjustment averaged across partner transitions 3 through 5 for all participants. Shaded area indicates 95% confidence intervals.

### Effects of Early Social Experiences Persist Through Development

Prior work has shown that the amount of time children spend in daycare during the first few years of life can influence their ability to adjust their communication to a partner at age 5, independent of familial influences^10^. To evaluate whether these early social experiences continue to affect stereotype- or interaction-based communicative adjustments at age 17, we conducted Spearman rank correlations between daycare attendance and communicative adjustment at each partner transition. A cluster-based statistical analysis, accounting for multiple comparisons across transitions, identified a cluster of negative correlations between daycare attendance and stereotype-driven adjustment spanning the third through the fifth transition (Fig. 2c). Put differently, individuals who spent more time in daycare exhibited reduced stereotype-driven adjustments during partner transitions 3 through 5 (Fig. 2d, *ρ*_(91)_ = -0.384, *p* = 0.001). As the timing of this effect coincided with a reduction in stereotype-driven adjustment observed across the whole group (Fig. 1c), we investigated whether individuals who spent more time in daycare also showed a faster rate of interaction-driven convergence in their communicative behavior toward the matched understanding of the two partners. A Spearman rank correlation of the slope of stereotype-driven adjustments over partner transitions found that this was the case (Fig. S5, *ρ*_(88)_ = -0.228, *p* = 0.031), suggesting a more rapid decline in stereotype contributions, influenced by the interaction dynamics, among individuals with higher exposure to daycare. Additional control analyses indicated that these interaction-dependent effects of daycare attendance were specific to communicative aspects of behavior and could not be explained by participants’ education levels or sensitivity to feedback (SI results). Furthermore, the effects of daycare attendance on adjustment remained statistically significant when controlling for the familial environment, including parents’ socio-economic status and the presence of siblings, and when accounting for social experiences later in life, including the number of friends at age 7, extracurricular activities at age 12, and time spent with friends age 14 (Fig. S6 and SI results). Additionally, as discussed below, the findings were largely robust to controlling for stereotype-driven variance indexed by variation in right ACCg volume and thickness (*ρ*_(67)_ = -0.196, *p*_one-tailed_ = 0.054; *ρ*_(67)_ = -0.226, *p*_one-tailed_ = 0.031). Together, these findings indicate that the effects of early social experiences persisted through development and continue to influence interaction-driven communicative adjustments.

## Discussion

By exploiting the human tendency to spontaneously adjust communicative behavior directed at children^50–53^, this study identifies neuroanatomical and developmental factors modulating how stereotype beliefs about an interaction partner are combined with evidence arising from social interactions. Participants were asked to communicate nonverbally with an adult and a child. In reality, a role-blind confederate performed both roles, such that the two presumed partners differed only in terms of participants’ stereotype beliefs regarding their cognitive abilities. Behaviorally, communicative adjustments driven by stereotype beliefs were transiently prominent at the onset of interactions, then gradually declined as interaction-based evidence accumulated against the presumption of differing communicative understanding between the two partners. Neuroanatomically, stereotype-based adjustments covaried (∼27%) with structural features of a distributed right-hemispheric cortical network, including the right anterior cingulate gyrus (ACCg). Developmentally, interaction-based adjustments were accounted for (∼14%) by individual variation in exposure to social interactions around the second year of life, over and above effects of familial environment and social experiences acquired later in life. A number of control analyses further qualify the specificity of these findings – e.g., stereotype-driven variance indexed by neuroanatomical variation in the right ACCg could not account for the interaction-dependent effects of early social experiences on communicative adjustment, and vice versa. Together, these findings indicate that stereotypes and interaction-based evidence provide complementary, yet neuroanatomically and developmentally dissociable, contributions to interpersonal communication.

Our findings provide a unified framework that reconciles previously disparate approaches in social and developmental psychology (Fig. 3). These approaches have focused on either stereotype-based generalizations for representing others, or on the construction of understanding from the interaction dynamics^16,17,54^. Here, we characterize the relative dynamics of stereotype- and interaction-based adjustments, making both types of adjustments experimentally visible on a single metric. Stereotypes provide initial boundaries on mutual understanding^55,56^, and this study highlights the cognitive work required to precisely translate that abstract knowledge into the situation at hand. Namely, stereotype-based adjustments could not be directly based on retrieving surface features of previous child-directed behaviors, such as tone of voice or simpler words, due to the novelty of the communicative setting. Put differently, participants’ stereotype-based adjustments likely required a degree of abstraction from experiences with or knowledge of young children. Stereotype-based adjustments could not be accounted for by generic priming effects either^57^, given that they were specifically bound to communicatively relevant aspects of participants’ interactive behaviors. In fact, the magnitude of the adjustments (∼30 ms on an average of 1150 ms spent in the object location) points to a finely controlled bias rather than an explicitly designed strategy. These observations suggest that stereotypes support the construction of functionally precise abstractions tailored to meet the situational demands of social interaction, rather than merely offering inflexible generalizations^58^.

**Figure 3.**
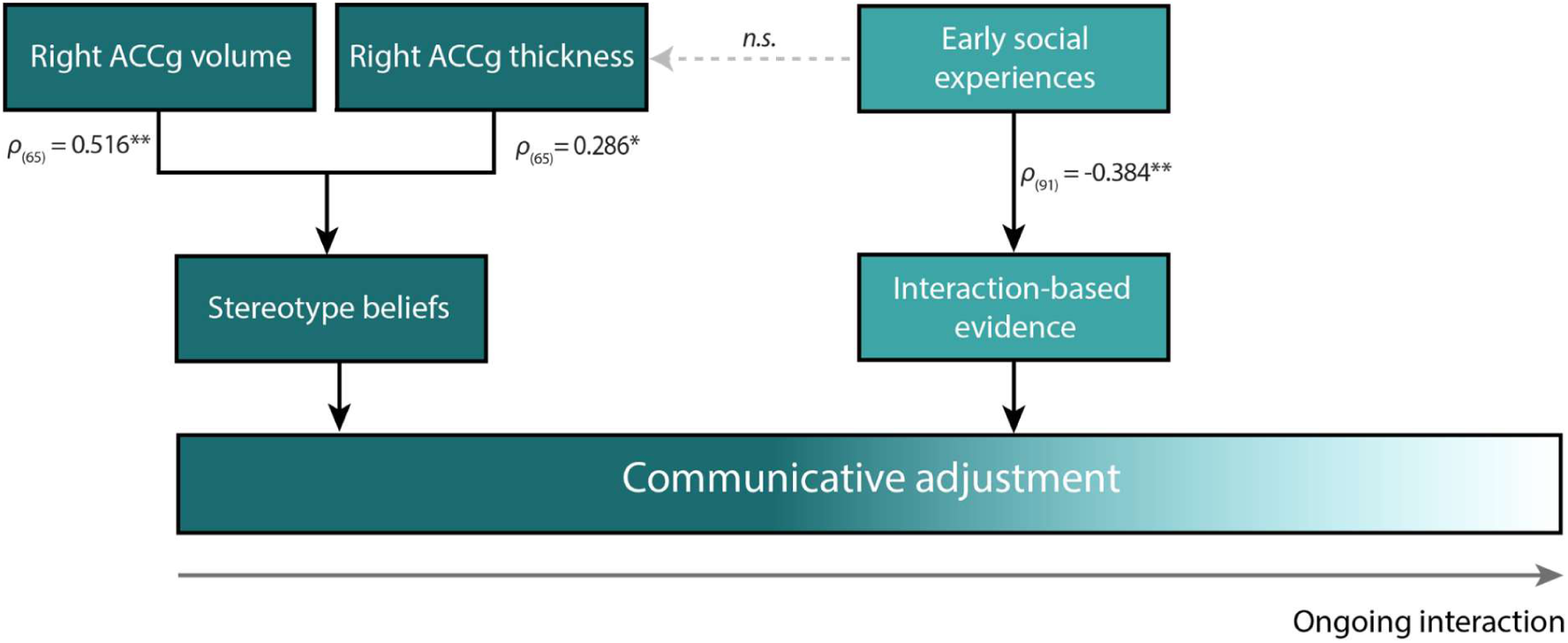
Summary model of the findings. The model depicts the dynamic interplay between stereotype beliefs and evidence derived from social interactions in shaping communicative adjustment toward an interaction partner. The impact of these factors varies based on an individual’s tendency to rely on stereotypes and integrate evidence from social interactions. These individual tendencies, in turn, are independently predicted by the structural integrity of the right anterior cingulate gyrus (ACCg) and early-life exposure to social interactions, collectively contributing to diverse rates at which individuals navigate and potentially overcome stereotype tendencies during social interactions.

The neuroanatomical findings of this study deepen the implications of these observations, establishing a structural basis for the notion that people differ in the application of stereotypes during social interactions^1^. The medial prefrontal cortex is known to facilitate mental state inferences based on stereotypical social knowledge^59–61^, possibly an instance of the broader function of this cortical region in supporting adaptive, schema-based behavior^62–64^. Notably, the anterior cingulate gyrus is instrumental in directing effort toward altruistically motivated behaviors^65–67^. This mechanism is likely crucial for the adaptation of preconceived notions into behaviors tailored to the present interactional context, such as communicating with greater emphasis to a child. The observed positive correlation between communicative adjustment and cortical thickness in the anterior cingulate gyrus may indicate a protracted, slower maturation of this cortical region in individuals exhibiting higher levels of stereotype usage. Pubertal brain maturation is marked by cortical thinning and synaptic pruning^68^, processes susceptible to acceleration by adverse early-life events^69^. Our finding that contemporary structural measures outperform brain scans acquired three years prior from the same individuals in predicting communicative adjustment suggests substantial developmental shifts in how social schemas are flexibly applied during adolescence. The observed increase in gray matter density and cortical thickness within this brain region might be a structural marker of this cognitive ability in interpersonal communication.

While stereotypes impact the way people approach a communicative interaction, some individuals are better than others at adjusting their reliance on these initial priors, making their communicative behaviors as informative as required given evidence derived from the ongoing interaction^70,71^. A major finding of this study is that interaction-based adjustments were predicted by participants’ time spent in daycare, a main source of exposure to interactions with unrelated conspecifics early in life. These early social experiences had long-lasting consequences on the ability to flexibly adjust communication to interaction partners. These consequences exceeded the effects of familial environment, a known influence on the emergence of false-belief understanding in early childhood^72–74^, as well as the effects of social experiences acquired later in life. This finding, combined with a closely related observation in five-year-olds^10^, suggests that the initial years of life may represent a sensitive developmental period for social interaction. We have previously reported that daycare experiences boost stereotype-based communicative adjustments in five-year-olds starting to master mentalizing abilities^10^. Here we show that, later in life, these same experiences boost interaction-based communicative adjustments in adolescents still refining their pragmatic abilities^75^. We suggest that the challenges of a daycare environment set infants and toddlers on a different socio-developmental trajectory. This trajectory might lead to an earlier acquisition of stereotype-based generalizations, and consequently a more fine-grained ability to manipulate and adjust these generalizations to individual minds later in development. The rationale here is that the social interactions afforded by a daycare environment offer a larger set of communicative challenges than those experienced within a relatively stereotyped familial environment. For instance, in daycare, a child needs to interact with rotating peers from different backgrounds and perspectives as well as ingratiate oneself with adult caregivers, all of which lack a genetic motive for collaborating and provisioning^76^.

In conclusion, this study demonstrates that humans can overcome stereotype tendencies within and through social interactions. The findings highlight the role played by the anterior cingulate gyrus in these tendencies while providing long-range longitudinal evidence that the ability to overcome stereotypes, based on evidence from social interactions, is sculpted by social experiences with unrelated conspecifics early in life. These findings offer a deeper understanding of how stereotypes and factual evidence are integrated in interpersonal communication, laying a foundation for future studies investigating systemic bias in society linked to stereotypes.

## Methods

### Participants

Participants were recruited from the Nijmegen Longitudinal Study on social-emotional development, which followed a community-based sample of 129 children and their families since infancy^77^. Ninety-six individuals from this longitudinal cohort agreed to participate shortly after their seventeenth birthday (47 females; 17.2 ± 0.2 y of age, M ± SEM). They were screened for a history of psychiatric and neurological problems and had normal or corrected-to-normal vision. Participants gave written informed consent according to institutional guidelines of the local ethics committee (Committee on Research Involving Human Subjects, Arnhem-Nijmegen region, The Netherlands) and received financial compensation for their participation. Seventy-two participants agreed to undergo MRI scanning for brain structural analysis. One participant experienced difficulties adhering to the task instructions and was excluded from further analysis. Another participant was excluded because of poor MRI scan quality, leaving seventy individuals for structural brain-behavior analysis. Additionally, participants agreed to provide a saliva sample for testing of testosterone levels as a proxy for pubertal maturation, which was used to account for individual differences in neurodevelopment^45^.

### Socio-Developmental Factors

Prior developmental work has shown that the amount of time children spend in daycare during the first few years of life can influence how flexibly they adjust their communication to a partner at age 5, over and above effects of familial environment^10^. In this study, we considered the same sources and indices of early developmental exposure to social interactions, obtained from the Nijmegen Longitudinal Study. Through a pre-registered analysis, we sought to determine if the consequences of these early-life social experiences persist through development to influence communicative abilities later in life^41^. Exposure to non-familial social interactions was indexed as the time spent in daycare, sampled at ages 15 and 28 months. This included regular daycare and childminders but not babysitting by biological kin including grandparents. We considered the average days per week across the two samples, after converting hours into days for the 28-month sample (1.4 ± 1.2 days/week). Familial environment was indexed with parents’ socio-economic status and the number of siblings. The former considered the average education level of both parents, sampled at 15 months of age (7-point scale: 5.2 ± 1.6 mean education level). Because parental education level and daycare attendance were correlated (*ρ*_(95)_ = 0.41, *p* < 0.001), parental education level was residualized against the working hours of the mother to avoid multicollinearity in the statistical models (correlation between parental education level and daycare attendance after correction for working hours of the mother: *ρ*_(88)_ = 0.18, *p* = 0.100). Number of siblings was sampled at 28 months of age. Because the greater majority of participants had one or no siblings (0.8 ± 0.6 siblings), we coded the presence of siblings as a dichotomous variable. We additionally considered participants’ exposure to social interactions later in life, also obtained from the Nijmegen Longitudinal Study. These late social experiences were indexed by the number of friends at age 7, extracurricular activities at age 12, and time spent with friends at age 14. Number of friends (6.01 ± 1.6 friends) and time spent with friends outside of school (10.86 ± 7.5 hours/week) were based on parent reports, whereas the measure of extracurricular activities was based on self-reports of how often children participated in sports, musical, theater, scouting, or other joint activities (5-point scale per item for a total of 25 points: 9.42 ± 2.47). Furthermore, we considered participants’ own education level at age 16 to control for individual differences in general learning ability (9-point scale: 5.4 ± 1.5).

### Task

We used the same two-player communication game as employed in previous experiments^8–11^. The game involves participant-partner dyads interacting on a digital game board with a 3×3 grid layout, visually presented on each player’s computer screen. On each of forty consecutive trials, their joint goal was for the partner (a confederate) to collect an acorn from a location on the game board shown to participants only (Fig. 1a). The acorn could be located in any of the thirteen white circles, and its trial-specific location was revealed to participants at the onset of the trial (phase 1). Given the experimental setup, participants could inform their partner by moving a bird avatar across the game board using a handheld game controller, visible to both players (phase 2). Four face buttons would move the bird horizontally or vertically to the center of each of the nine squares of the grid. There were no restrictions on planning time, but participants had only 5 seconds for moving the bird, as counted from the onset of the first move. Additionally, participants needed to return the bird to its starting position on the nest in the center of the grid. After 5 seconds, or earlier if participants pressed the stop button, a squirrel avatar would appear in the center, visible to both players and signaling the partner’s turn. Using a digital mouse, the partner could move the squirrel to the circle deemed to contain the acorn given the bird’s movements, selecting it by clicking the left mouse button (phase 3). Unlike the bird, there were no temporal or spatial restrictions on the movements of the squirrel on the game board. The squirrel movements were also visible to both players, allowing participants to observe and interpret how their movements of the bird were understood by their partner, in addition to explicit feedback on the successful retrieval of the acorn at the end of the trial (phase 4). Successful trials, in which participants had successfully guided the partner to the acorn and returned the bird to the nest, resulted in the presentation of a large acorn on the screen. A red cross was presented over a small acorn for unsuccessful trials. To challenge participants toward the generation of mutually understood communicative behaviors, task difficulty was manipulated across trials. This was achieved by introducing a disparity between the positions allowed to the bird and the possible locations of the acorn, i.e. the white circles. Namely, the bird could not be overlaid on the precise location of the acorn when a square contained more than one circle. Importantly, even in trials where the acorn was in a square with one circle, the participant still needed to make clear to the partner which square contained the acorn, disambiguating it from other squares visited by the bird while moving across the game board. Trials where the acorn was in a square with one, two, or three circles were pseudo-randomly intermixed such that there was an overall increase in difficulty over the course of the experiment.

### Experimental Design

During the experiment, participants sat in an experimental room facing a monitor displaying the digital game board. A confederate partner sat in another room facing a monitor that displayed the same game board. First, each participant was familiarized with the experimental setup (3 trials). During these familiarization trials, the participant was encouraged to move the bird around the game board, experiencing the constraints on the bird’s movements as described in the task section. The participant was also instructed to return the bird to the nest in the center of the grid at the end of their movements. This task requirement was reinforced by a continuous sound of a bird chirping that started when the participant moved away from the nest and stopped upon returning the bird to the nest. Next, each participant was informed that they would be playing the game online with two other players in turns, an adult and a five-year-old child (Fig. 1A). They were told that the game partners were sitting in other rooms and could see the bird and the digital game board on their monitors. In reality, the confederate performed the roles of both partners while remaining blind to which role he was performing in any given trial. The confederate did not know the location of the acorn and could infer that location only from the movements of the participant. Participants alternately interacted with the presumed child and adult partner over eight blocks of five trials, for a total of forty trials. A digital photograph of the current partner was presented in full screen before each block and in the top-right corner during each block. There were two presentation orders of adult-child partners and two sets of acorn configurations, counterbalanced over participants. The game was programmed and presented using Presentation software on a Windows XP personal computer (Neurobehavioral Systems, Albany, CA, USA). To prevent the leaking of information about the manipulation of the study, there was no structured debriefing following the experiment. However, participants were offered the opportunity to comment on the experiment. None did so.

### Stereotype-Based Adjustments

Past research using the same task has repeatedly shown that participants spontaneously spend longer time with the bird in the square containing the acorn when interacting with a presumed younger partner^8–11^, a behavior functionally equivalent to the prosodic modifications observed in verbal communication directed at children^51^. This study exploited these communicative adjustments as a real-time quantitative marker of stereotype beliefs about an interaction partner, measured in milliseconds. Paired-samples *t*-tests were used to test for differences in the average time spent in the location of the acorn as a function of the presumed partner. To account for generic changes in movement speed related to task difficulty or procedural learning effects, we controlled for the number of candidate locations in squares containing the acorn as well as planning time and time spent in other visited locations of the game board. This was achieved by residualizing time spent in the acorn location against these control variables using a linear mixed-effect model implemented in R’s lme4 package^78^. Trials with missing values for one or more control variables were excluded from this analysis (16% of all trials). To capture the temporal dynamics of stereotype-driven communicative adjustment, we calculated the difference in average time spent in the acorn location between the eight consecutive experimental blocks of five trials, producing adjustment estimates for seven partner transitions (see experimental structure in Fig. 1c). The sign of these partner transitions considered the expected direction of the adjustment, i.e. communicative adjustments were multiplied by +1 and -1 for adult-child and child-adult transitions respectively. The resulting adjustment estimates were entered into a cluster-based statistical analysis correcting for multiple comparisons across transitions, using one-sample *t*-tests with clusters defined using a threshold-free cluster enhancement (TCFE) of *p* < 0.05 based on 10,000 Monte Carlo permutations^42^.

### Interaction-Based Adjustments

Communicative adjustments driven by the interaction dynamics were quantified through convergence in behavior directed at the presumed child and adult partner. This measure considered the same dependent variable as stereotype-based adjustments, namely the time spent in the location of the acorn when moving across the game board. Spearman rank correlations were used to test for a relationship between early exposure to social interactions and stereotype-based adjustment at each partner transition. As above, the resulting correlation estimates were entered into a cluster-based statistical analysis correcting for multiple comparisons across transitions. To assess the dynamic nature of interaction-driven adjustments, we considered the linear slope of stereotype-based adjustment over partner transitions, starting from its statistical onset at the second partner transition and extending through the fifth transition, the final transition shaped by early life social experiences. This slope captured the rate of communicative convergence toward the matched understanding of the two partners in each individual. The slopes were entered into a partial Spearman rank correlation analysis evaluating the effects of early-life exposure to social interactions on the rate of communicative convergence, controlling for the intercept which was highly correlated with the slope (*ρ*_(91)_ = -0.834 *p* < .001). Follow-up analyses were conducted using partial Spearman rank correlations to test whether the relationship between early social experiences and communicative convergence remains statistically significant after controlling for effects of familial environment and social experiences acquired later in life. Finally, we used partial Spearman rank correlations to test whether the interaction-dependent effects of daycare attendance on communicative adjustment could be accounted for by stereotype-driven variance indexed by individual variation in ACCg volume and thickness.

### Brain Structural Measures

Anatomical images for structural brain-behavior analysis were acquired using 3T Siemens Trio and Prisma scanners. Both scanners used a 32-channel head coil and MP-RAGE sequence (TR/TE, 2.30 s/3.03 ms; voxel size, 1 x 1 x 1 mm^3^; FOV, 256 mm), and produced comparable image quality, as determined using SPM12’s computational anatomy toolbox, CAT12. We considered two in-vivo measures of neuroanatomical structure, based on voxel- and surface-based morphometry of gray matter density and cortical thickness, respectively. These measures were obtained by segmenting the anatomical images into gray matter, white matter, and cerebrospinal fluid, and then coregistering them to MNI space using DARTEL. For voxel-based analysis, the gray matter segments were smoothed with an isotropic 8 mm full-width half-maximum (FWHM) Gaussian kernel. To account for individual differences in brain hemispheric volume, gray matter densities were expressed as a function of gray matter content in homotopic areas of the opposite hemisphere, producing a voxelwise index of gray matter asymmetry^43^. For surface-based analysis, cortical surfaces were reconstructed from the gray matter segments and smoothed with a 15 mm FWHM kernel^44^. The resulting voxel- and surface-based measures were used as predictors of communicative adjustment in separate multiple linear regression analyses, controlling for gender, scanner type, and salivary testosterone levels. Follow-up analyses included Spearman rank correlations between communicative adjustment and ACCg volume and thickness, as determined using statistical masking (*p*_FWE_ < 0.05, family-wise error corrected) and the Desikan-Killiany anatomical atlas (caudal anterior cingulate)^46^. Additionally, generalizability of the structural MRI findings was assessed in two additional datasets, including the same sample at age 14 and an independent sample performing the same task.

### Salivary Testosterone Levels

Salivary testosterone levels were measured in order to control for individual differences in pubertal maturation in the brain structural analysis. Prior research has indicated that testosterone levels in both boys and girls correlate with pubertal status, as determined by physical examination and self-reported puberty ratings^45,79,80^. Participants provided two saliva samples, collected 2 hours apart after 10:00 AM, in order to minimize the effects of diurnal fluctuations in endogenous testosterone levels. The samples (∼2 ml) were collected using Salicap (IBL) containers, stored at -24°C, and later analyzed using a competitive chemiluminescence immunoassay (CLIA) with a sensitivity of 0.0025 ng/ml. Participants were instructed to not eat, smoke, or drink (except water) at least one hour before the start of the experiment. The values of the first collected sample were log-transformed and standardized within each sex before being used as covariates in the brain structural analysis^45^.

## Acknowledgements

The authors would like to thank all families that participated in the Nijmegen Longitudinal Study. This work was supported by the NWO Gravitation Grant (024.001.006) to the Language in Interaction Consortium. IT is supported by an Advanced Grant from the European Research Council (101054559), KR by a Consolidator Grant from the European Research Council (ERC_CoG_772337) and HCMN by a NWO Research Talent Grant (406-13-022). The NLS neuroimaging waves were supported by a starting grant from the European Research Council (ERC_StG2012_313749 awarded to KR). The data collection at age 16 was supported by a NWO Research Talent Grant (406-12-110) from the Netherlands Organization for Scientific Research awarded to J. Loes Pouwels.

## Author contributions

SBJK data analysis and interpretation, wrote the original draft. AT study design, data collection. HCMN study design, data collection. AHNC project conception, study design. KR project conception, study design. JB data interpretation. IT study design, data interpretation, wrote the original draft. AS study design, data interpretation, wrote the original draft. All authors contributed to the interpretation and preparation of the final manuscript.

## Data availability

The datasets generated and analyzed during the current study will be made available in the Donders Repository (https://data.donders.ru.nl).

## Code availability

The underlying code for this study will also be made available in the Donders Repository.

## Supplementary Information

### SI Results

In the following sections, we report a number of additional analyses designed to verify the specificity and robustness of the findings reported in the main text.

### Communicative specificity of the stereotype-based adjustments

We performed additional analyses to verify whether the stereotype-driven adjustments reported in the main analysis (Fig. 1B) were specific to communicatively relevant portions of the game board and did not influence other aspects of participants’ behavior such as planning time and time spent in other locations of the game board. We report the results of Bayesian linear mixed models, considering planning time and time spent in other locations as dependent variables, presumed partner (adult, child) as a within-subject factor, and participant as a random factor to account for individual differences. We used the *brms* package in R with default priors for all models^81^. We considered effects to be statistically significant in case the 95% credible interval (CI) of the posterior distributions did not contain zero. Additionally, posterior probability (*pp*) values are reported, which can be interpreted as traditional *p*-values at an alpha level of 0.025^82^. The models found that the dependent variables of participants’ planning time (B = -0.01, 95% CI = [-0.03 0.01], *pp* = 0.143) and time spent in other locations of the game board (Fig. S1; B = - 0.01, 95% CI = [-0.03 0.01], *pp* = 0.191) were matched between both presumed partners. Moreover, a cluster-based analysis found no communicative adjustments in other locations of the game board across any partner transition. These results support the notion that the stereotype-driven adjustments were specific to communicatively relevant aspects of participant behavior.

### Consistency of the confederate’s task performance

We performed additional analyses to determine if the confederate showed consistent behavior and understanding in the roles of the child and adult partner. We report the results of Bayesian (generalized) linear mixed models using the same specifications as above, but with the proportion of correctly solved trials and the response time of the confederate as dependent variables. The results of the analysis showed that the proportion of correctly solved trials (Fig. S2a; B = 0.09, 95% CI = [-0.03 0.22], *pp* = 0.075) and the confederate’s response time (Fig. S2c; B = 0.01, 95% CI = [-0.01 0.04], *pp* = 0.124) were matched between both partners. An additional analysis focused on the consistency of the confederate’s performance across participants found that there was no statistically significant relationship between the order in which the participants were tested and the proportion of correctly solved trials (Fig. S2b; *ρ*_(95)_ = 0.08, *p* = 0.420) or confederate’s response time (Fig. S2d; *ρ*_(95)_ = 0.10, *p* = 0.340). These results suggest that the confederate’s task performance was consistent within and across participants.

### Generalizability of the structural brain-behavior findings

We performed additional analyses to assess the generalizability of the identified association between the right anterior cingulate gyrus (ACCg) and stereotype-driven communicative adjustment (Fig. 2a). We used two additional datasets, including the same sample at age 14 (45 individuals; 23 females) and an independent sample that performed the same task (27 individuals; 27 males; 22.67 ± 3.41 y of age). For the participants at age 14, identical systems and protocols were used to acquire the T1-weighted MRI data. For the independent sample, structural images were acquired with a 1.5T Siemens Avanto scanner using a single-shot MPRAGE sequence (TR/TE, 1.73 s/2.95 ms; voxel size, 1 x 1 x 1 mm^3^; FOV, 256 mm). This dataset comprised the placebo group from a study on the effects of intranasal oxytocin administration on communicative adjustment^8,83^. We applied the same voxel-based morphometry pipeline as used in the main analysis to both datasets, and extracted right ACCg volume using a statistical mask based on the findings reported in the main text. For the participants at age 14, we correlated right ACCg volume with communicative adjustment at age 17. For the independent sample, we correlated right ACCg volume with communicative adjustment during partner transition 6 through 8, when adjustment was statistically prominent^83^. As shown in Fig. S4, Spearman rank correlations found that communicative adjustment was positively associated with volume in the right ACCg in both the same sample at age 14 (*ρ*_(45)_ = 0.376, *p* = 0.011) and the independent sample (*ρ*_(27)_ = 0.389, *p* = 0.045). Moreover, even with a strong correlation between right ACCg volume at age 14 and the corresponding volume at age 17 (*ρ*_(42)_ = 0.887, *p* < 0.001), the association with stereotype-driven adjustment at age 17 remained significant after adjusting for right ACCg volume at age 14 (*ρ*_(36)_ = 0.389, *p* = 0.016). These results suggest the timeliness and replicability of the structural brain-behavior relationships highlighted in the main analysis across various datasets.

### Specificity and persistence of the effects of early social experience

We conducted additional analyses to ascertain whether the effects of early social experiences on communicative adjustment (Fig. 2c) retained significance after accounting for the effects of familial environment (parents’ socio-economic status and the presence of siblings), late social experiences (number of friends at age 7, extracurricular activities at age 12, and time spent with friends at age 14), and participants’ education levels (at age 16). We used partial Spearman rank correlation analyzes to control for variance accounted for by these social environmental factors and found that the inverse relationship between daycare attendance and communicative adjustment remained statistically significant. Moreover, daycare attendance did not covary with any of the late social experiences (all *ρ* ≤ 0.169), and as seen in Fig. S6, was the strongest predictor of communicative adjustment at age 17. These results suggest that the effects of early social experiences on communicative adjustment are specific and persistent.

### Communicative specificity of the interaction-based adjustments

We performed additional analyses to verify whether the effects of early social experiences on communicative adjustment were specific to communicatively relevant portions of the game board and did not influence other aspects of participant behavior, including planning time and time spent in other locations of the game board. We report the results of Bayesian linear mixed models, considering planning time and time spent in other locations as dependent variables, presumed partner (adult, child) as a within-subject factor, participant as a random factor, and daycare attendance and familial environment as predictors. We also modeled the interaction between these predictors and the factor of partner. We found that daycare attendance was positively associated with planning time (B = 0.10, 95% CI = [0.01 0.19], *pp* = 0.017), but that this relationship was not modulated by partner (B = -0.01, 95% CI = -0.03 0.01, *pp* = 0.313). Daycare attendance was not associated with other indices of task performance such as proportion of correctly solved trials (B = -0.06, 95% CI = [-0.35 0.23], *pp* = 0.337) and time spent in other locations of the game board (B = 0.11, 95% CI = [-0.01 0.24], *pp* = 0.037). Additionally, time spent in daycare did not modulate the relationship between partner and the proportion of correctly solved trials (B = -0.00, 95% CI = [-0.12 0.11], *pp* = 0.480) or time spent in other visited locations (B = 0.00 95% CI = [-0.01 0.02], *pp* = 0.285). Moreover, a cluster-based analysis found no significant association between daycare attendance and communicative adjustments in other locations across any partner transition, nor was the rate of convergence in time spent in these other locations significantly associated with daycare attendance (*ρ*_(91)_ = 0.018, *p* = 0.862). Finally, we investigated whether the effects of early social experiences on communicative adjustment could be explained by varying sensitivities to feedback among participants. A Bayesian generalized linear mixed model considering success as the dependent variable and daycare attendance, trial number, and presumed partner as predictors found no statistically significant relationship between presumed partners and the dynamics of success (B = -0.00, 95% CI = [-0.02 0.01], *pp* = 0.341). Moreover, there was no statistically significant interaction between daycare attendance, trial number, and partner (B = -0.00, 95% CI = [-0.01 0.01], *pp* = 0.311). Together, these results suggest that the effects of daycare attendance on communicative adjustment were specific to communicatively relevant portions of the game board and not influenced by other factors.

### SI Figures

**Fig. S1.**
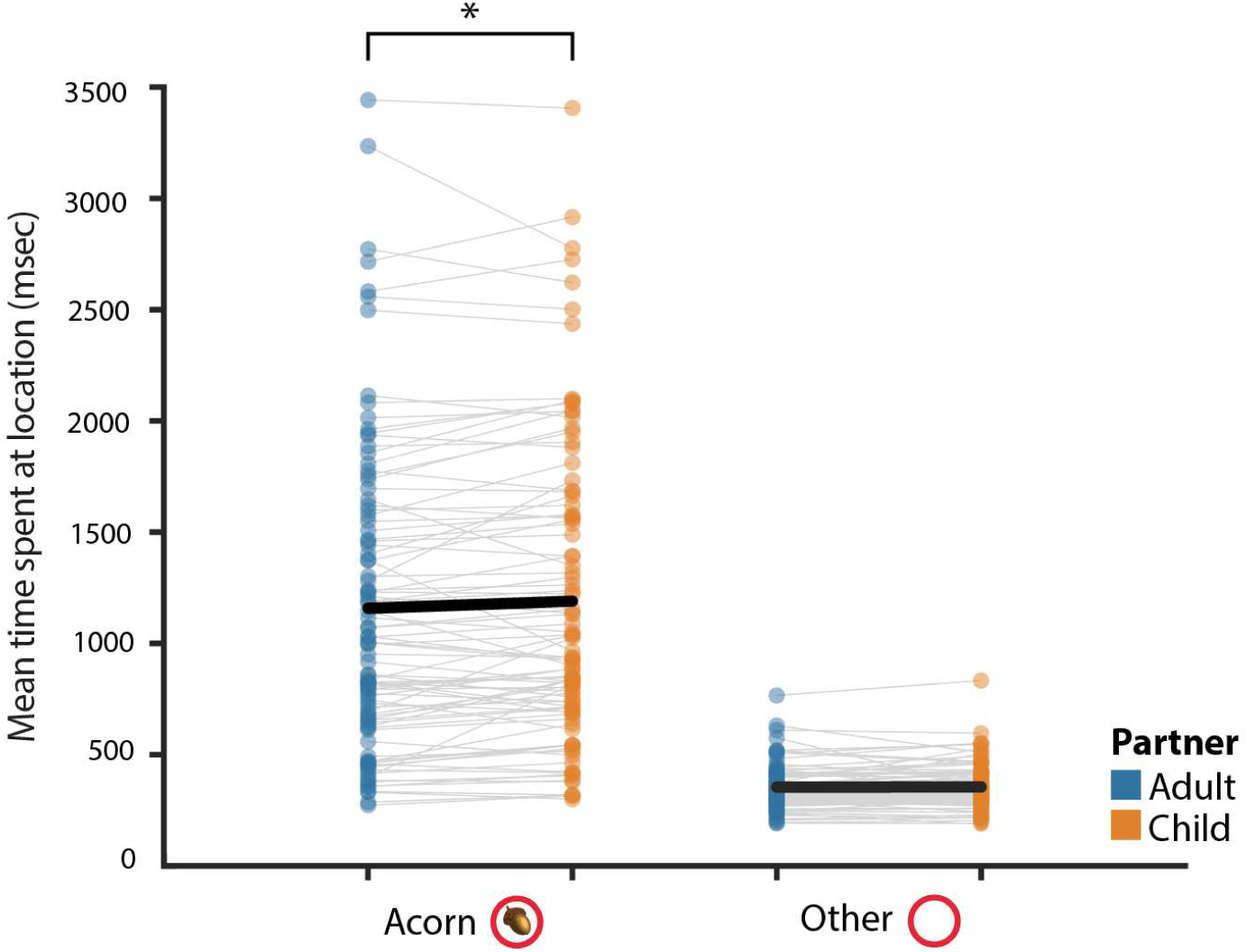
Average time spent in different locations of the game board as a function of the presumed partner. Participants spent more time in the location of the acorn when interacting with the child partner compared to the adult partner (*M difference* = 31.58 ms, *t*_(94)_ = 2.31, *p* = 0.023, Cohen’s *d* = 0.24). There was no significant difference in the time spent in other locations between the two presumed partners (*M difference* = -3.46 ms, *t*_(94)_ = -0.69, *p* = 0.491, Cohen’s *d* = 0.07). Dots in the graph represent individual data points and the black lines indicate averages.

**Fig. S2.**
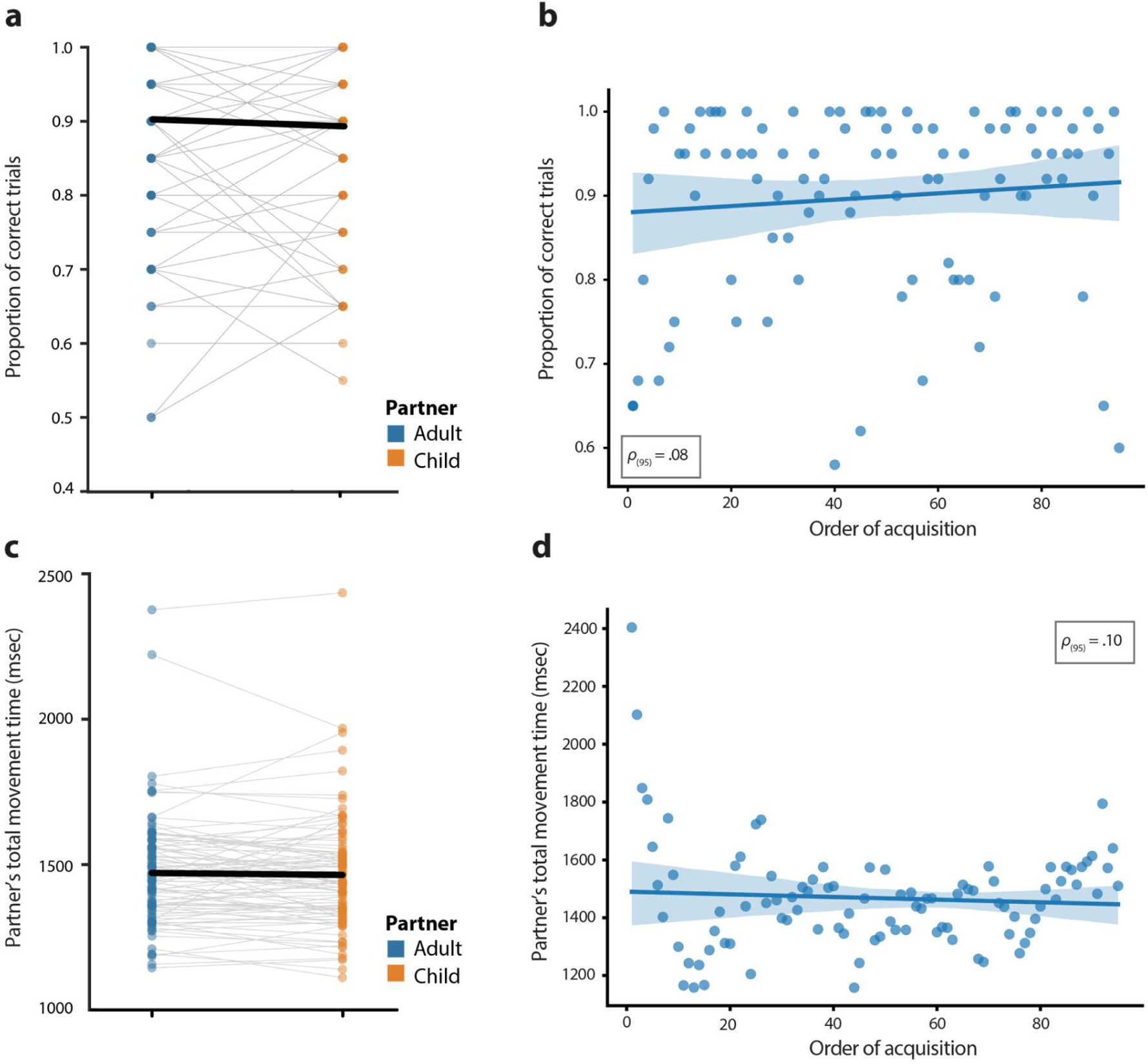
Consistency of the confederate’s task performance both within and across participants. (**a**) The proportion of correctly solved trials was consistent between the adult and child partner within each participant, and (**b**) this consistency was maintained across participants and did not change as a function of the order in which participants were tested. (**c**) The confederate’s response time was consistent between both partners within each participant, and (**d**) this consistency did not change as a function of the acquisition order.

**Fig. S3.**
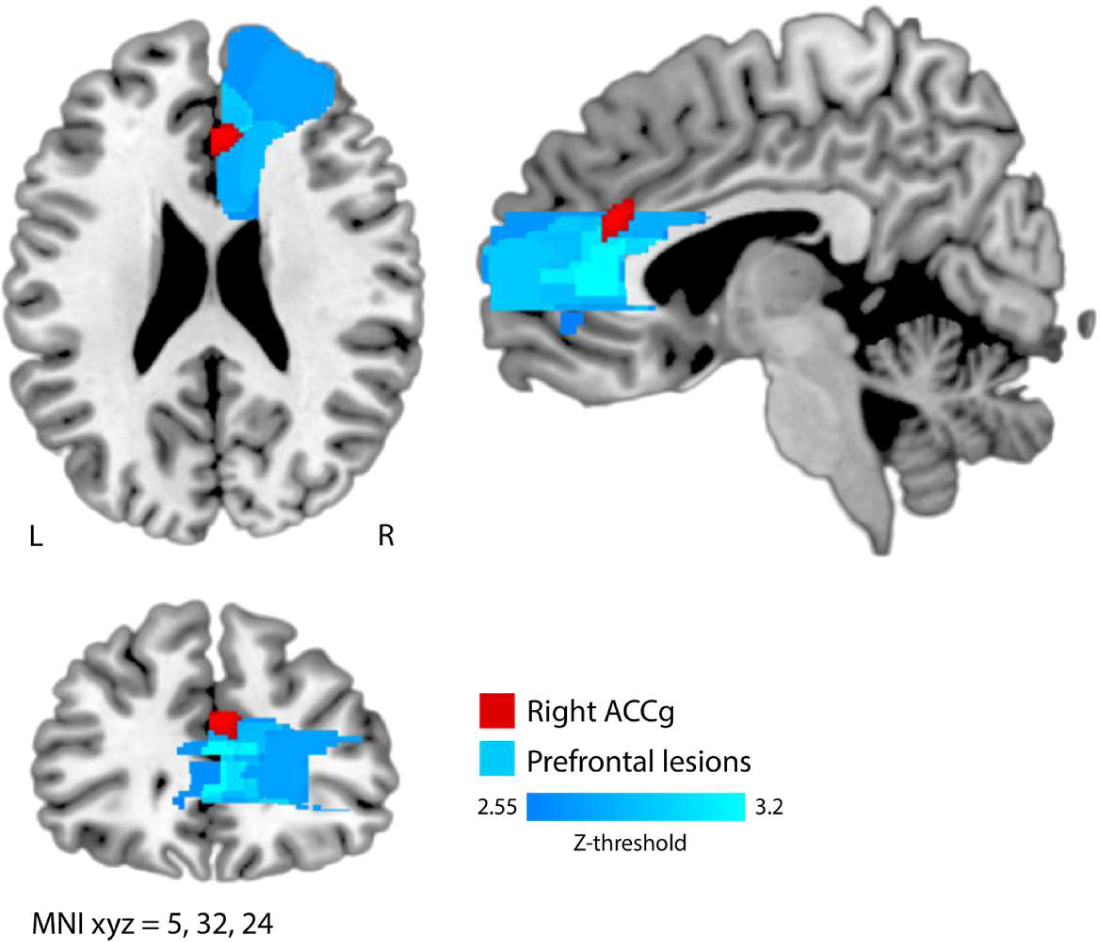
Anatomical overlay of the cluster in the anterior cingulate gyrus (ACCg) found to be associated with stereotype-driven communicative adjustment (red, from Fig. 2a) on a lesion-based voxelwise map of prefrontal brain regions that are crucial for adjusting communication based on stereotype beliefs (blue, adapted from^11^). Z-values correspond to a significance level of 0.05 > *p* > 0.001 (two-tailed).

**Fig. S4.**
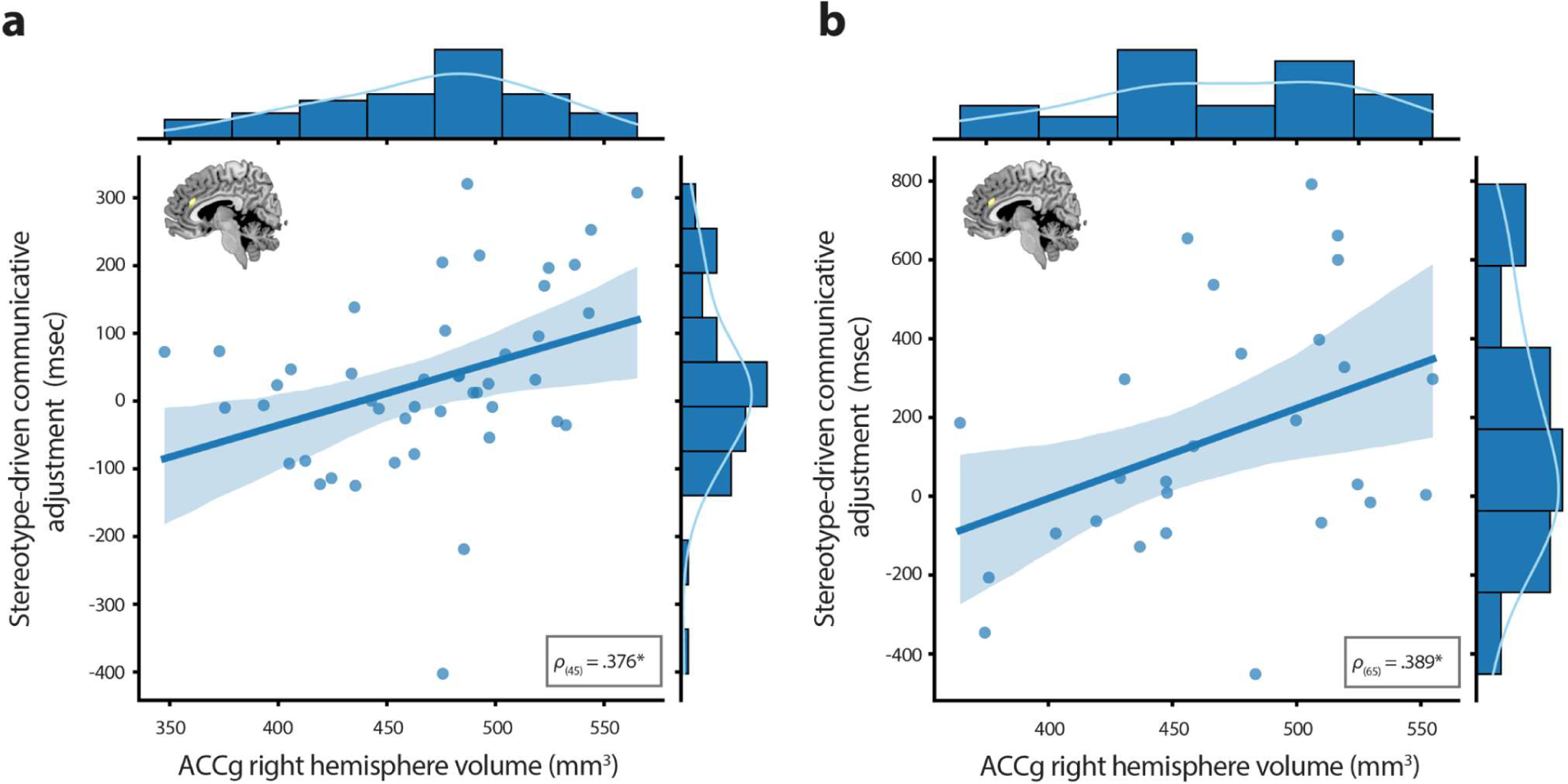
Replication of the association between gray matter density in the anterior cingulate gyrus (ACCg) and communicative adjustment to stereotype beliefs in two additional datasets, namely (**a**) the same participants at age 14 and (**b**) an independent sample performing the same task.

**Fig. S5.**
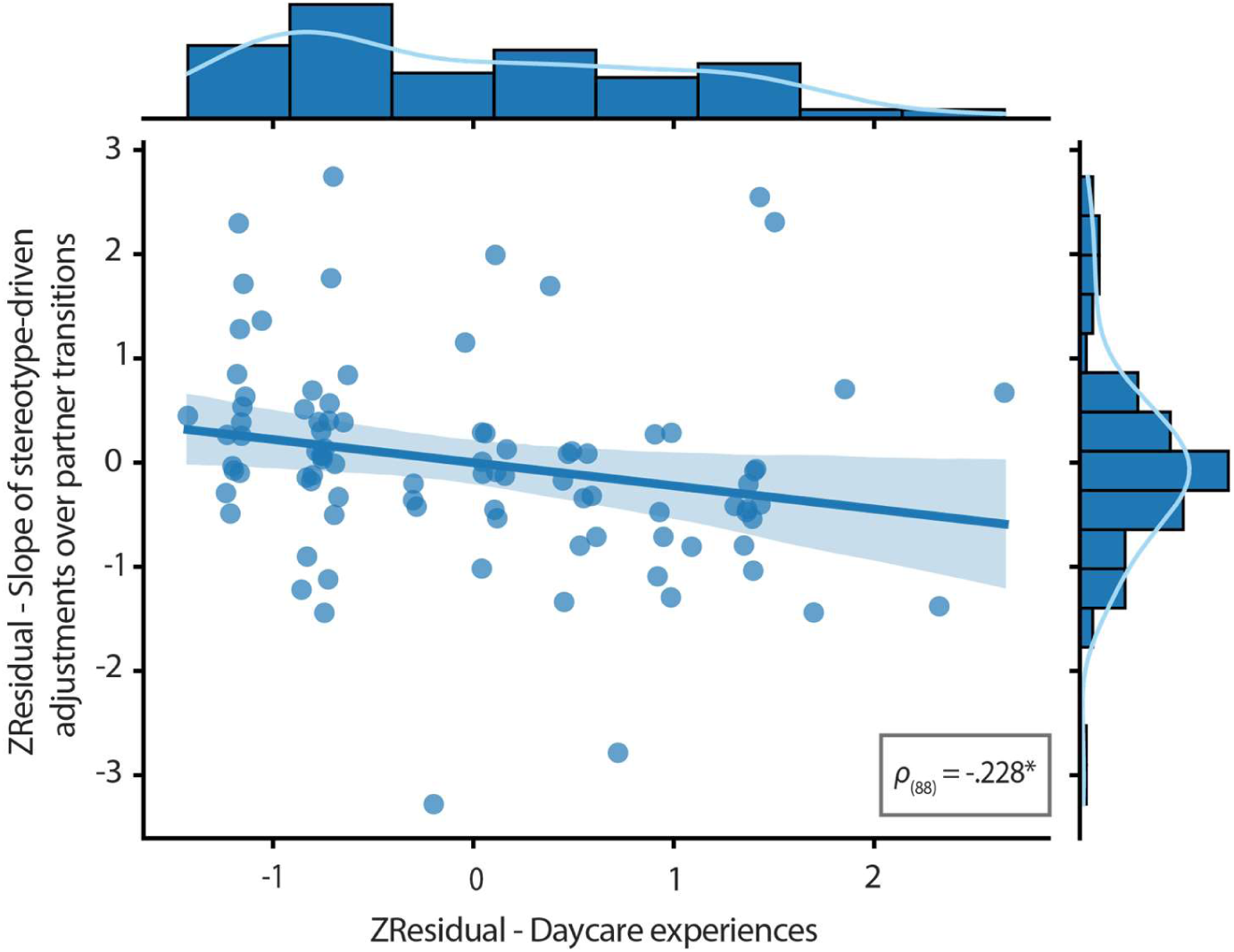
Impact of early social experiences on the rate of communicative convergence at age 17. Time spent in daycare was predictive of the slope’s steepness derived from the temporal dynamics of (declining) stereotype-driven communicative adjustment. ZResidual = standardized residual, corrected for the intercept of the linear function fitted through partner transitions.

**Fig. S6.**
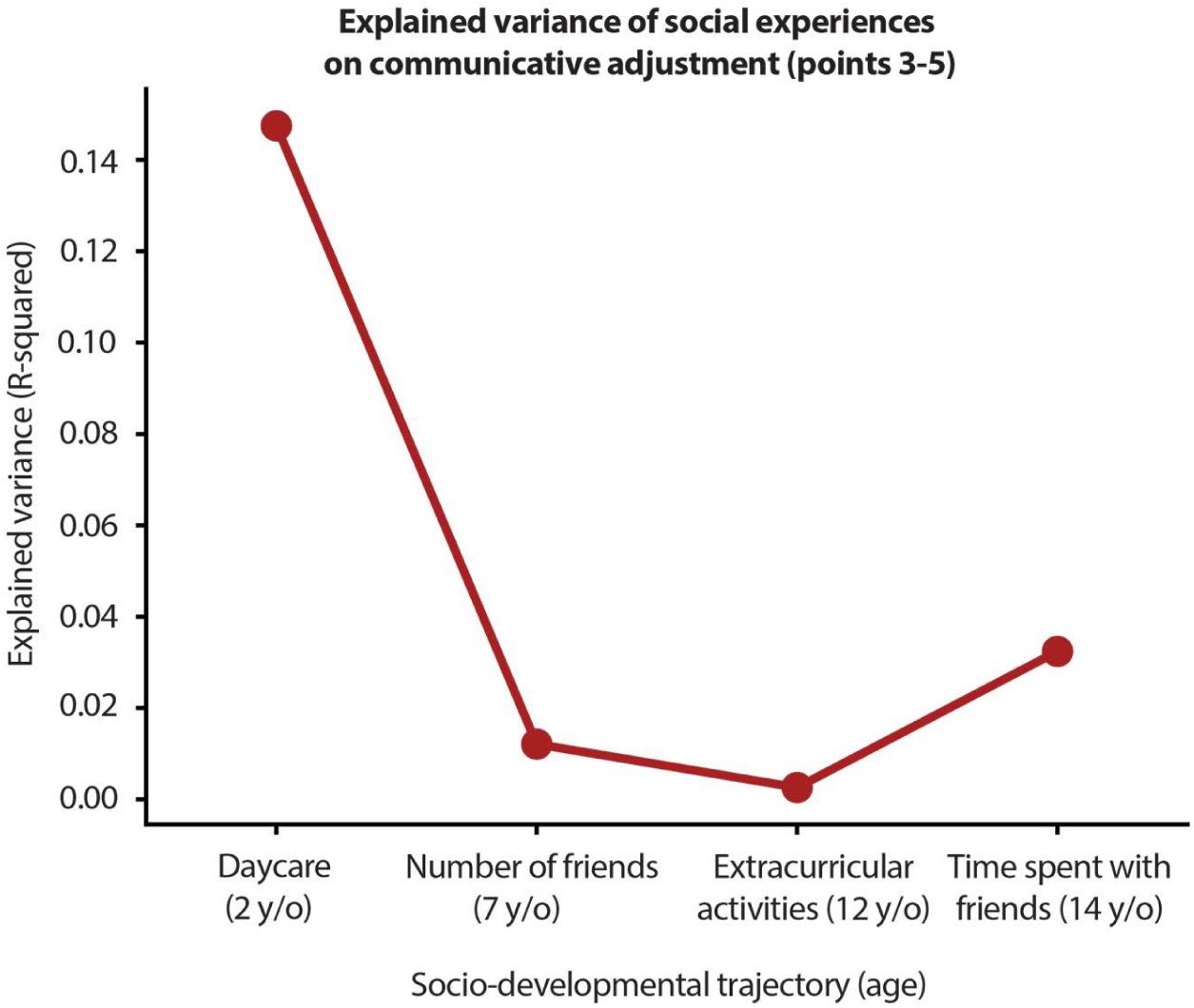
Effects of early and late social experiences on interaction-based communicative adjustment at age 17. Time spent in daycare around the second year of life accounted for over 14% of the variance in (reduced) stereotype-driven communicative adjustment. By contrast, late social experiences (number of friends at age 7, extracurricular activities at age 12, and time spent with friends at age 14) did not significantly contribute to the variance in interaction-based communicative adjustment.

